# *Deinbollia onanae* (Sapindaceae), a new, Endangered, montane tree species from the Cameroon Highlands

**DOI:** 10.1101/2020.10.24.353680

**Authors:** Martin Cheek, Jean Michel Onana

## Abstract

*Deinbollia onanae* (Sapindaceae-Litchi clade) is here formally named and characterised as a new species to science, previously known as *Deinbollia sp. 2*. Cameroon has the highest species-diversity and species endemism known in this African-Western Indian Ocean genus of 42 species. *Deinbollia onanae* is an infrequent tree species known from five locations in surviving islands of montane (or upper submontane) forest along the line of the Cameroon Highlands. It is here assessed as Endangered according to the IUCN 2012 standard, threatened mainly by clearance of forest for agriculture. The majority of tree species characteristic of montane forest (above 2000 m alt.) in the Cameroon Highlands are also widespread in East African mountains (i.e. are Afromontane). *Deinbollia onanae* is one of only a very small number of species that are endemic (globally restricted to) the mountain range. It is postulated that this new species is in a sister relationship with *Deinbollia oreophila,* which is a frequent species of a lower (submontane) altitudinal band of the same range. It is further postulated that seed dispersal is or was by frugivorous birds, potentially turacos, alternatively by primates such as Preuss s monkey.

## INTRODUCTION

As part of the project to designate Important Plant Areas (IPAs) in Cameroon (also known as Tropical Important Plant Areas or TIPAs), we are striving to name, assess the conservation status and include in IPAs (*Darbyshire et al., 2017*) rare and threatened plant species in the surviving, threatened natural habitat of the Cross-Sanaga interval (*Cheek et al., 2001*). Several of these species were previously designated as new to science but not formally published in a series of checklists (see below) ranging over much of the Cross-Sanaga interval. The Cross-Sanaga has the highest vascular plant species diversity per degree square in tropical Africa (*Barthlott et al., 1996*) but natural habitat is being steadily being cleared, predominantly for agriculture.

In this paper we formally describe and name as *Deinbollia onanae* Cheek a high-altitude tree species formerly designated as “*Deinbollia* sp. 2” (*Harvey et al., 2004, Cheek et al., 2004, Cheek et al., 2010*).

The genus *Deinbollia* Schum. & Thonn. is traditionally place in the tribe Sapindeae DC. and is characterised by its 1-pinnate, imparipinnate leaves, flowers with petals well developed and about the same in number as the imbricate sepals, the petals with a well-developed ligule (or appendage) on the adaxial surface and with stamens 9–30 in number, the intrastaminal disc central, the edge with more than 5 shallow ridges. The fruits develop 1–3 indehiscent, apocarpous fleshy mericarps (*Fouilloy & Hallé, 1973*).

Molecular Phylogenetic sampling of Sapindaceae is incomplete with many African genera not represented, as can be seen in *Buerki et al.* (*2009*). In that study *Deinbollia* is represented by six samples of four species, all from Madagascar (on which limited basis it appears monophyletic) and is resolved in the informally named ‘Litchi Group’ of genera, where it is in a sister relationship to a subclade comprising the genera *Lepisanthes* Blume (Africa to Asia) *Atalaya* Blume (American) and *Pseudima* Radlk. (American) (*Buerki et al., 2009*). The delimitation of Sapindaceae in this paper follows the evidence of *Buerki et al.* (*2010*), that is, excluding Aceraceae, Hippocastanaceae and Xanthoceraceae which have sometimes been included within it.

*Deinbollia* has 41 accepted species, one shared between Africa, Reunion and Madagascar, 5 endemic to Madagascar, and 35 species restricted to subsaharan continental Africa. The species predominantly occur in lowland evergreen forest and are absent from countries that lack this habitat such as Rwanda, Burundi, Swaziland and Lesotho (high altitude) and Namibia, Botswana, Eritrea, Mali and Burkina Faso (low rainfall and lacking significant evergreen forest). The highest species diversity is found in Cameroon, with 16 species (Plants of the World Online accessed May 2020). Cameroon has the highest levels of country-level endemism in the genus. Ten of the Cameroon species were assessed as globally threatened with extinction (Cheek in *Onana & Cheek 2011: 314– 316*). In contrast only 10 species are recorded for the whole of West Tropical Africa (*Keay, 1958*). Since the Flore Du Cameroun account was published (*Fouilloy & Hallé, 1973*), several further species apart from those listed below, were published for Cameroon by *Thomas* (*1986*). The genus was last revised by *Radlkofer* (*1932*).

In the 21^st^ century only two new species to science have been published in the genus, *Deinbollia mezilii* D.W.Thomas & D.J.Harris (*Thomas & Harris, 2000*) and *D. oreophila* Cheek (*Cheek & Etuge 2009*), both from Cameroon. But specimens often remain unidentified in herbaria. For example, 16 specimens unidentified to species are listed in the Gabon Checklist (*Sosef et al.,2005*). The genus has no major uses but the fruits of several species are reported as being edible by humans, and the seeds are probably primate-dispersed or dispersed by large frugivorous birds, and the flowers probably bee-pollinated (*Cheek & Etuge, 2009*), while the bark of *D. grandifolius* Hook.f. is used medicinally and the wood for planks (*Burkill, 2006*).

## METHODS & MATERIALS

Fieldwork in Cameroon resulting in the specimens cited in this paper was conducted under the terms of the series of Memoranda of Collaboration between IRAD (Institute for Agronomic Research and Development)-National Herbarium of Cameroon and Royal Botanic Gardens, Kew beginning in 1992, the most recent of which is valid until 5^th^ Sept. 2021. The most recent research permit issued for fieldwork under these agreements was 000146/MINRESI/B00/C00/C10/C12 (issued 28 Nov 2019), and the export permit number was 098/IRAD/DG/CRRA-NK/SSRB/12/2019 (issued 19 Dec 2019). At the Royal Botanic Gardens, Fieldwork was approved by the Institutional Review Board of Kew entitled the Overseas Fieldwork Committee (OFC) for which the most recent registration number was OFC 807-3 (2019). The most complete set of duplicates for all specimens made was deposited at YA, the remainder exported to K for identification and distribution following standard practice. Field work methodology followed was *Cheek & Cable* (*1997*).

Herbarium citations follow Index Herbariorum (*Thiers et al., 2020*). Specimens indicated “!” were seen by one or more of the authors, and were studied at K, P, WAG, and YA. The National Herbarium of Cameroon, YA, was also searched for additional material of the new taxon as was Tropicos (http://legacy.tropicos.org/SpecimenSearch.aspx). During the time that this paper was researched in 2019–2020, it was not possible to obtain physical access to material at WAG (due to the transfer of WAG to Naturalis, Leiden, subsequent construction work, and covid-19 travel and access restrictions). However images for WAG specimens were studied at https://bioportal.naturalis.nl/?language=en and those from P at https://science.mnhn.fr/institution/mnhn/collection/p/item/search/form?lang=en_US. We also searched *JStor Global Plants* (*2020*) for additional type material of the genus not already represented at K.

Binomial authorities follow the International Plant Names Index (*IPNI, 2020*). The conservation assessment was made using the categories and criteria of *IUCN* (*2012*). GeoCAT was used to calculate red list metrics (*Bachman et al., 2011*). Herbarium material was examined with a Leica Wild M8 dissecting binocular microscope fitted with an eyepiece graticule measuring in units of 0.025 mm at maximum magnification. The drawing was made with the same equipment using Leica 308700 camera lucida attachment. Flowers from herbarium specimens of the new species described below were soaked in warm water to rehydrate the flowers, allowing dissection, characterisation and measurement. The terms and format of the description follow the conventions of (*Cheek & Etuge, 2009*).

## RESULTS

### TAXONOMIC TREATMENT

*Deinbollia* sp. 2, because it has leaves less than 1 m long, only sparsely hairy on the lower surface, leaflets more than 15 cm long and sepals adaxially glabrous, flower buds very sparsely hairy and less than 5 mm diam. borne on a branched inflorescence 10–30 cm long, keys out in the Flore Du Cameroun treatment of *Deinbollia* (*Fouilloy & Hallé, 1973*) to a couplet leading to *D. grandifolia* Hook.f. and *D. maxima* Gilg. However, it differs from these two species in having (2–)8–11-jugate (not 4–7-jugate), and in other characters shown in table 1.

**Table 1.** Characters separating *Deinbollia onanae* from *D. grandifolius* and *D. maxima.* Characters for the first two species taken from *Fouilloy & Hallé* (*1973*).

|  | *Deinbollia grandifolia* | *Deinbollia maxima* | *Deinbollia onanae* |
| --- | --- | --- | --- |
| Leaves | (5–)7-jugate | 4–6-jugate | (2–)8–11-jugate |
| Leaflet width | 5–8 cm | 6–8(–10) cm | (2.1–)2.5–4 cm |
| N°s pair of secondary nerves (distal leaflets) | 12–14(–16) | 8 – 10 | (12–)17–18 |

The affinities of *Deinbollia* sp. 2 may instead be with, however, the recently described *D. oreophila* since this species also occurs at altitude in the Cameroon Highlands and both species share numerous raised lenticels and also leaflets with high length: breadth ratios and with high numbers of secondary nerves. In fact, at two locations, Mt Kupe and Bali Ngemba, the two species are sympatric and their altitudinal ranges can overlap slightly (*Cheek et al., 2004, Harvey et al., 2004*). As the only two species of the genus to grow at altitude in the Cameroon Highlands, there is a possibility that they might be confused with each other. The two species can be separated using table 2.

**Table 2.** The more significant differences between *Deinbollia onanae* and *Deinbollia oreophila*.

|  | *Deinbollia oreophila* | *Deinbollia onanae* |
| --- | --- | --- |
| Height at maturing | 0.8–3(–5) m | (4–)5–10(–15) m |
| Stem indumentum | Glabrous | Simple hairy, sparse to dense, glabrescent. |
| Lenticels | Highly conspicuous, bright white, contrasting with epidermis | Inconspicuous, grey-brown, concolorous with epidermis |
| Length of leaves (flowering stems) | 25–63 cm | (14–)60–70 cm |
| Number of leaflets per leaf (flowering stems) | (4–)6–8(–10) | (4–)16–23 |
| Width of leaflets (flowering stems). | (3–)5.5–9(–10.2) cm | (2.1–)2.5–4 cm |
| N° secondary nerves each side of midrib | (7–)9–14(–17) | (15–)17–18 |
| Indumentum of lower surface of leaf blade | Glabrous | Inconspicuously simple hairy |
| Sepals | Orbicular, margins glabrous | Ovate, margins hairy |
| Petals | Oblong or obovate, base cuneate; adaxial appendage surface glabrous | Rhombic or spatulate, basal claw (stalk); adaxial appendage surface hairy |
| Staminal filaments of male flowers | Proximal half glabrous. | Entire length densely hairy. |
| Ovary of female flowers | 3-lobed, surface with very sparse, stout hairs | 2-lobed, densely hair with fine hairs |
| Altitudinal range | (880–)1000–2050 m | (1200–)2000–2200 m |

### Deinbollia onanae Cheek *sp. nov.* – Fig. 1

Similar to but differing from *Deinbollia oreophila* Cheek in the length of leaves of flowering stems (14–)60–70 cm, number of leaflets per leaves (4–)16–23, width of leaflets (2.1–)2.5–4 cm, number of secondary nerves on each side of midrib (15–)17–18, not respectively, 25–63 cm, (4–)6–8(–10), (2.1–)2.5–4 cm and (15–)17–18; stems with lenticels brown, concolorous and inconspicuous, not discolorous, bright white and conspicuous; ovary bilocular not trilocular. Type: *Etuge* 3600 (holotype K000337729!; isotypes MO!, WAG0336084!, WAG0336083!, YA0057050!), Cameroon, *Mt Oku and the Ijim Ridge*, Aboh to Tum fl. 22 Nov. 1996.

*Deinbollia cf. pinnata* Schum. & Thonn. sensu Cheek, p 162 in *Cheek et al.,* (*2000*). *Deinbollia* sp. 2 sensu Cheek in *Harvey et al.,* (*2004**: 125***); Cheek & Etuge in *Cheek et al.,* (*2004*: 399); Cheek in *Cheek et al.,*(*2010*: 143, fig 23).

Monoecious tree or treelet (4–)5–10(–15) m tall, when in flower, lacking exudate or scent when wounded, sparingly branched, nearly glabrous, apart from the inflorescence. Stems of flowering branches terete 1–1.5 cm diameter, solid (not hollow), second internode below apical inflorescence 2–2.5 cm long, outer epidermis pale grey-brown, contrasting with the darker brown bases of the adjoining petiolar pulvini, lenticels dense, raised, elliptic, 0.6–1.1 mm long, concolorous, inconspicuous, glabrescent, hairs sparse to dense, dark brown, cylindric 0.1–0.5 mm long.

Leaves alternate, pinnately compound, (14–)60–70 cm long; leaflets (4–)16–23 per leaf on flowering stems, leaflets 10–14 per leaf on leaves of juvenile trees. Petiole (4–)9.5–20.8 cm long, terete, c. 4 mm diameter at midpoint, drying pale yellow; basal pulvini dark brown; rhachis (4.5–)32–44 cm long, (2–)8–11-jugate on flowering stems, 5–7-jugate on non-flowering stems of juvenile trees, the upper surface of the distal half strongly convex with two lateral wings, glabrescent with sparse inconspicuous hairs (*de Wilde* 4555), or with dense dark brown appressed hairs (*Cable* 3386). Leaflets mostly oblong (6.6–)14–19.5 × (2.1–)2.5–4 cm, (but leaflets of sterile branches to 6.5 cm wide), acumen c. 1 cm long, base broadly acute, slightly asymmetric, (basalmost leaflets lanceolate and about half the length of the other leaflets) lateral nerves and midrib yellow, raised above and below, convex, (15–)17–18 on each side of the midrib, nearly brochidodromous, the lateral nerve apices forming a weak irregular submarginal nerve, stronger branches uniting with the secondary nerve above, intersecondary nerves strong, parallel to the secondaries, tertiary and quaternary nerves reticulate raised yellow and conspicuous, on both surfaces, contrasting with the pale grey-green areolae (except in *Cable* 3386(K) where they are concolorous and so inconspicuous above, possibly an artefact of poor drying); upper surface glabrous, lower surface with inconspicuous, minute, cylindrical, glossy dark-brown hairs c. 0.25 mm long, distributed very sparsely along the midrib and secondary nerves, absent from mature leaves of non-flowering specimens (e.g. *Cheek* 8709) but then the same hair type present on axillary buds and young leaves; petiolules yellow, 2–5 mm long.

Inflorescence a 80–120-flowered, loose, terminal panicle 25 × 10 cm; auxiliary inflorescences sometimes present in the axils of the distal 1–4 leaves (*Cheek* 13625); peduncle of terminal inflorescences 0–2 cm long; rhachis internodes (1–)2–3 cm long, shortest in the distal portion; first order bracts caducous; indumentum brown hairy; primary branches 10–20 per inflorescence, 2–8 cm long, each bearing (1–)2–5 partial-inflorescences; partial-peduncles 0–5 mm long, apex with a cluster of 3–5 bracteoles; bracteoles subulate to narrowly lanceolate, 2–3 mm long, apex narrowly acute, partial-inflorescences (1–)3-flowered in glomerules, pedicels erect, terete, 3–4 × 1.5 mm (female), 4–5 × 1mm (male), sparsely puberulent, hairs 0.1–0.5 mm long.

Flowers white, scent not recorded, flower buds c. 4 mm diam., open flowers c. 6 × 7 mm. Calyx with sepals 5(–6), orbicular to broadly ovate, concave, green colour, 4–5 × 3.5–4.5 mm apex obtuse. Corolla apex slightly exserted from calyx, petals rhombic or spatulate. Male flowers (Fig. 1C). Petals 5(–6), white, rhombic c. 5 ×x 3 mm, apex obtuse-acute, base cuneate, margins densely ciliate, hairs 0.3 mm long, outer surface glabrous, inner surface glabrous in distal half, proximal half compressed funneliform with ventral appendage adnate at margins, retuse (notched) for 0.5 mm at midline, adaxial surface moderately densely hairy, hairs c. 0.3 mm long. Extra-staminal disc torus-like, glabrous, irregular, outer wall convex, lacking constrictions or teeth with c. 15 poorly defined lobes, 2.5–3 mm wide, c. 0.8 mm high. Stamens c. 15, erect, slightly exserted by 1–2 mm at anthesis, c. 5–6.5 mm long; filament 4–5 mm long, straight, densely puberulent the entire length (Fig. 1D); anthers yellow, ovate-ellipsoid, 1–1.3 mm long. Ovary (vestigial, Fig. 1E) bilobed, c. 1 × 1.5 mm densely appressed hairy, hairs c. 0.5 mm; style 0.7 mm long, glabrous.

**Fig.1.**
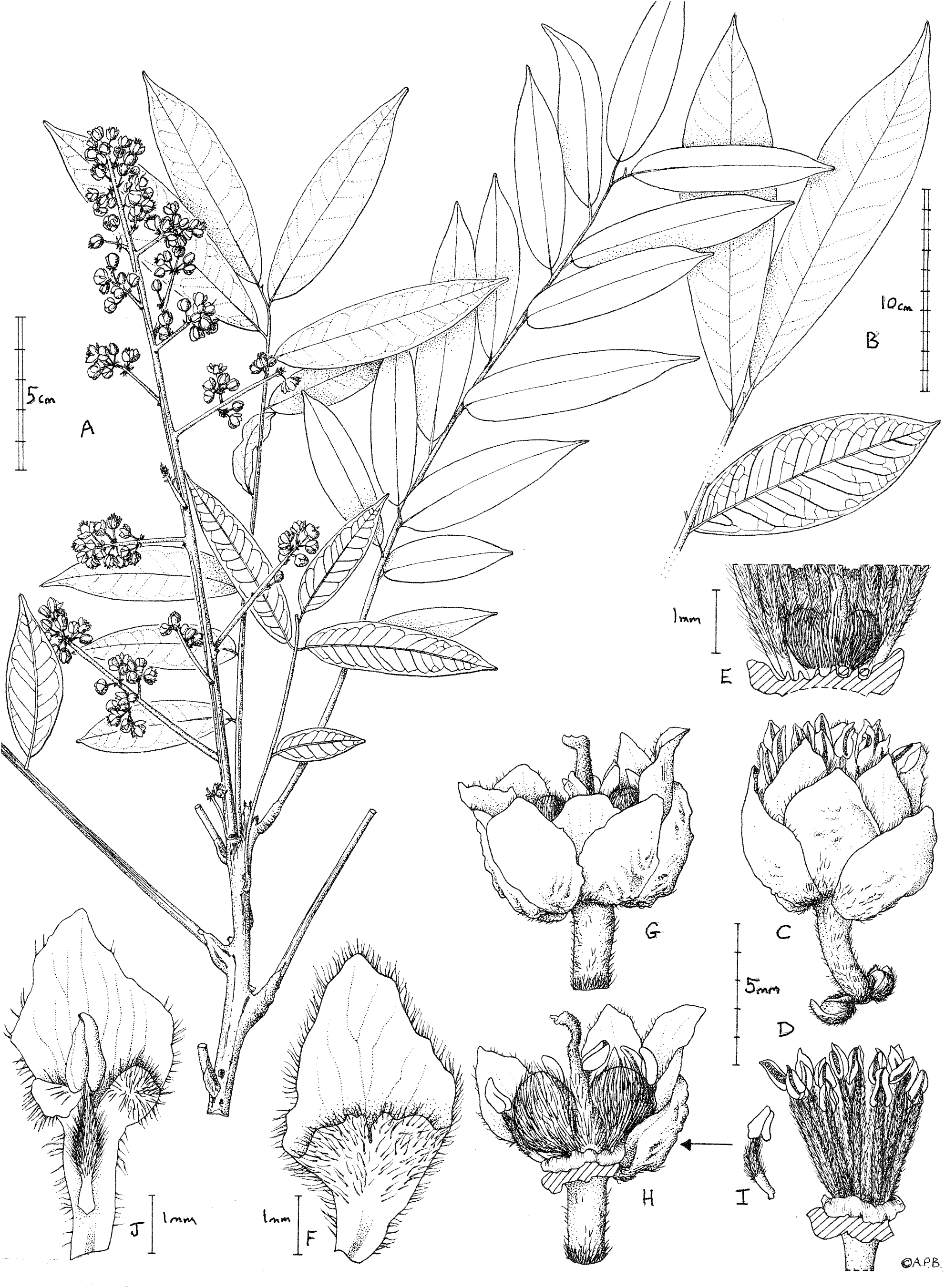
*Deinbollia onanae.* (A) habit, flowering branch; (B) detail from a large leaf showing apex and distal leaves (adaxial surfaces) and second leaf from the base (abaxial surface); (C) male flower lateral view; (D) male flower, petals and sepals removed to show the extra staminal disc and androecium; (E) base of D (male flower) showing the vestigial gynoecium and disc cut to show notches holding filament bases; (F) petal, adaxial surface, male flower; (G) female flower, lateral view; (H) female flower (with 3 sepals, 2 petals and anterior stamens removed) to show gynoecium and disc; (I) stamen from female flower; (J)petal, adaxial surface, of female flower with stamen. **A, C-J** from *de Wilde et al.* 4553 (K); **B** from *Cheek* 13436 (K). Drawn by Andrew Brown.

Female flowers (Fig. 1G), with sepals and petals as the male flowers, but petals c. 6 × 2.6–2.9 mm, usually detaching with a stamen attached, probably due to interlocking hairs (see Fig. 1 J), proximal two-thirds claw-like, c. 0.7 mm wide, margin sparsely and irregularly ciliate; ventral appendage with apex deeply bilobed, lobes c. 1 mm × 1 mm; disc as in male flower. Stamens c. 10 (see Fig. 1I), included at anthesis, filament c. 2.5 mm long, proximal half to quarter glabrous, distal part densely hairy; anther as male flowers but indehiscent; ovary bilobed (see Fig. 1 H), 3.2 × 5 mm, indumentum as male flower, style c. 5 mm long, apical 1 mm, curved, surface papillate-minutely puberulent, apex subcapitate. Infructescence, mature fruit and seed unknown.

### Phenology

Flowering in November-December; fruiting probably in August-September.

### Local name and uses

none are known. “Onana’s *Deinbollia* is suggested as a common name.

### Etymology

The specific epithet of *Deinbollia onanae* means ‘of Onana’ commemorating Dr Jean-Michel Onana, currently Lecturer in Botany at University of Yaoundé I, Cameroon, champion of plant conservation in Cameroon, specialist in Sapindales (Burseraceae, author of Flore Du Cameroun Burseraceae (*Onana, 2017*), co-chair of the IUCN Central African Red List Authority for Plants, former Head of the National Herbarium of Cameroon (2005–2016), co-author of the Red Data Book of the Plants of Cameroon (*Onana & Cheek, 2011*) and the Taxonomic Checklist of the Vascular Plants of Cameroon (*Onana, 2011*). He led field teams of YA staff working with those of K that resulted in the collection of several of the specimens of this species and personally collected this species in the field (*Onana* 1600, K, YA).

### Distribution & ecology

known only from the Cameroon Highlands of Cameroon (not yet recorded from Nigeria). Upper submontane & montane evergreen forest, sometimes in gallery forest; (1200-)2050–2200 m alt.

### Additional specimens: CAMEROON

**South West Region**, *Mt Kupe*, near main summit, immature fr., 26 June 1996, *Cable* 3386 (K000197863!, YA!); **North West Region.** *Bali Ngemba Forest Reserve*, April 2002, *Onana* 1600 (YA!); *Mt Oku and the Ijim Ridge*: above Laikom, st. 21 Nov..1996, *Cheek* 8709 (K000337728! YA!); *Dom*, Kinjinjang Rock, st. 25 Sept. 2006, *Cheek* 13436 (K000580433!; YA!); ibid. Forest Patch 1, fl. buds, 27 Sept. 2006, *Cheek* 13625 (K000580434!, MO!,US!, YA!); ibid., Javelong Forest, st. 29 April 2005, *Pollard* 1400 (K000580432!; YA!); **Adamaoua Region**, c. 120 km E of Ngaoundéré, 15 km NE of Belel, falls in Koudini River, alt. ± 1200 m, fl. 4 Dec. 1964, *W.J.J.O. & J.J.F.E. de Wilde, B.E.E. de Wilde-Duyfjes* 4555 (K000593309!; K000593310!, WAG1269760!, YA)

### Notes

*Deinbollia onanae* first came to our attention in 2000 when completing the “Plants of Kilum-Ijim” (*Cheek et al., 2000*). Two specimens of *Deinbollia* matched no other and were named *Deinbollia cf. pinnata* (*Cheek et al., 2000*). In subsequent surveys this taxon was more explicitly referred to as a new species: *Deinbollia* sp. 2 (*Harvey et al., 2004, Cheek et al., 2004, Cheek et al., 2009*). However, the earliest known collection was made in 1964 (*W.J.J.O. & J.J.F.E. de Wilde, de Wilde-Duyfjes* 4555(K)).

### Conservation

*Deinbollia onanae* is rare at each of its five known locations so far as is known. Despite many thousands of herbarium specimens being collected at Kilum-Ijim, at Mt Kupe and the Bakossi Mts, and at Bali Ngemba (*Cheek et al., 2000; Cheek et al., 2004; Harvey et al., 2006*) only two specimens of this species at two sites, were made at each of the first two locations and only one at the third location. Surveys at other sites in the Cameroon Highlands and elsewhere, e.g at Mt Cameroon and at the Lebialem Highlands, failed to find this species (*Cheek et al., 1996; Cable & Cheek 1998; Harvey et al., 2010; Cheek et al., 2011*). However, at Dom, where a targetted search for this species was made by the first author, three specimens were made, each representing single, isolated trees (*Cheek et al., 2010*). No more individuals than these were found. At Adamaoua it has only been collected once, and only a single tree was then noted (*W.J.J.O. & J.J.F.E. de Wilde, B.E.E. de Wilde-Duyfjes* 4555(K)). None of these locations is formally protected for nature conservation. Tree cutting for timber and habitat clearance for agriculture has long been known to be a threat at all but the last of these locations (references cited above). We assess the area of occupancy of *Deinbollia onanae* as 28 km² using the IUCN preferred 4 km² cell size. Therefore, we assess this species as Endangered, EN B2ab(iii) using the *IUCN* (*2012*) standard. We suggest that this species be included in forest restoration plantings within its natural range to partly reverse its move to extinction. However, the likely large (c. 1 cm diam.), thin-walled seeds are probably recalcitrant, so not suitable for conventional seed-banking, and should not be allowed to be dried before sowing.

## DISCUSSION

The discovery of a threatened, new species to science from surviving natural habitat in the Cameroon Highlands is not unusual. At most of the five locations from which we here describe *Deinbollia onanae,* additional new or resurrected species to science, all threatened with extinction, have been documented in recent years. At Mt Kupe for example, *Coffea montekupensis* Stoffelen (*Stoffelen et al., 1996*) and more recently the new species and genus to science *Kupeantha kupensis* Cheek & Sonké (*Cheek et al., 2018a*). At Bali Ngemba, *Leptonychia kamerunensis* Engler & K. Krause (*Cheek et al., 2013*), *Psychotria babatwoensis* Cheek (*Cheek et al., 2009*) and *Allophylus ujori* Cheek (*Cheek & Etuge, 2009b*), at Mt Oku and the Ijim Ridge *Kniphofia reflexa* Marais (*Maisels et al., 2000*), *Scleria cheekii* Bauters (*Bauters et al.*, 2018), while at Dom, the endemic epiphytic sedge *Coleochloa domensis* Musaya & D.A Simpson (*Muasya et al., 2010*). No additional such species are known from the Adamaoua location, probably because it less completely sampled than the preceeding four.

However, *Deinbollia onanae* is exceptional among these aforementioned species in that it is a new species of tree of predominantly montane forest. The many other newly discovered for science, resurrected or rediscovered plant species of the Cameroon Highlands have either been herbs or shrubs or are derived from submontane habitats (800–2000 m altitude). The division between montane and submontane forest is well-marked in Cameroon (*Cheek et al., 1996; Cheek et al., 2000*), although some species of tree, like *Deinbollia onanae* can occur on either side of the 2000 m contour. The tree species diversity of the montane forest is low compared with adjoining submontane forest, and in contrast, with very few Cameroon Highland endemic tree species. Almost all montane tree species of the Cameroon Highlands are widespread in montane forest in Africa (Afromontane) occurring also east of the Congo Basin in the rift mountains of East Africa. Apart from *Deinbollia onanae,* the only other montane tree species endemic to the Cameroon Highland chain are the nearly extinct *Ternstroemia cameroonensis* Cheek (*Cheek et al., 2017*) and the more common and widespread *Schleffera mannii* (Hook.f.)Harms (*Keay, 1958).*

The high altitudinal range of *Deinbollia onanae* is unrivalled west of the Congo basin by any other species of the genus. Elsewhere in Africa it is matched only by *Deinbollia kilimandscharica* Taub., of mountains from Ethiopia to Malawi, reported to achieve 2250 m elevation in Tanzania (*Davies & Verdcourt, 1998*). Most species of the genus in tropical Africa are lowland forest shrubs, in the Cameroon Highlands only *Deinbollia oreophila* also occurs regularly at altitude, and is largely confined to the submontane forest band being recorded from (880–)1000–1900(–2050) m altitude where it is often relatively frequent (*Cheek & Etuge, 2009*). We postulate based on their shared morphological characters that these two may be sister species that have segregated between two adjacent altitudinally based vegetation types in a similar way to certain clades of bird species in the Cameroon Highlands such as the Turaco (*Njabo & Sorensen, 2009*). This hypothesis needs testing. It would most readily done by a comprehensive species-level molecular phylogenomic study of *Deinbollia* as has been achieved in several other genera, such as *Nepenthes* L.f. (*Murphy et al., 2020*).

The fruits of *Deinbollia onanae* are expected to be similar to those of other species of the genus, i.e., fleshy, indehiscent and large-seeded, suggesting that the now intermittent distribution of this species, along a line c. 570 km along peaks of the Cameroon Highland line, is due to dispersal by animals. Dispersal is or was, possibly by a fruit-eating bird such as Bannerman’s Turaco (*Tauraco bannermani*), a species restricted to forest above 2200 m altitude (*Njabo & Sorensen 2009*). Alternatively, dispersal might be by a primate species such as Preuss’s monkey (*Allochrocebus preussi*), which lives in high altitude forest up to 2500 m altitude. Both species are threatened with extinction and have been assessed as Endangered under the IUCN standard. Formerly the range of *Deinbollia onanae* may have once been more continuous along the mountain range than today, but it was likely greatly reduced when forest was cleared for agriculture. Large sections of the range, such as the Bamenda Highlands, are now so denuded of the original forest that they are today referred to as “The grasslands”. It has been estimated in such areas that over 96.5% of the original forest has been lost (*Cheek et al., 2000*).

## CONCLUSIONS

Such discoveries as *Deinbollia onanae* underline the urgency for publishing further discoveries while it is still possible since threats to such newly discovered for science species are clear and current, putting these species at high risk of extinction. About 2000 new species of vascular plant have been discovered each year for the last decade or more (*Cheek et al., 2020*). Until species are known to science, they cannot be assessed for their conservation status and the possibility of protecting them is reduced (*Cheek et al., 2020*). Documented extinctions of plant species are increasing, e.g. *Oxygyne triandra* Schltr. of Cameroon is now known to be globally extinct (*Cheek et al., 2018b*). In some cases species appear to be extinct even before they are known to science, such as *Vepris bali* Cheek, once sympatric with *Deinbollia onanae* at Bali Ngemba (*Cheek et al., 2018c*), and elsewhere, *Nepenthes maximoides* Cheek (*King & Cheek, 2020*). Most of the >800 Cameroonian species in the Red Data Book for the plants of Cameroon are threatened with extinction due to habitat clearance or degradation, especially of forest for small-holder and plantation agriculture e.g. oil palm, following logging (*Onana & Cheek, 2011*). Efforts are now being made to delimit the highest priority areas in Cameroon for plant conservation as Tropical Important Plant Areas (TIPAs) using the revised IPA criteria set out in *Darbyshire et al.,* (*2017*). This is intended to help avoid the global extinction of additional endemic species such as the Endangered *Deinbollia onanae* which will be included in the proposed IPA s of Mt Kupe, Bali Ngemba, Kilum-Ijim and Dom.

## Acknowledgements

This paper was completed as part of the Cameroon Tropical Important Plant Areas Project, supported by the Players of Peoples Postcode Lottery. The second author’s contribution to this paper was made possible by visits from Cameroon to RBG, Kew, U.K. sponsored by the Bentham-Moxon Trust of RBG, Kew. Most of the specimens cited in this paper were collected with the support of volunteers of Earthwatch Europe, Oxford and by our colleagues Kenneth Tah, Olivier Sene, Victor Nana, Verina Ingram, David Okebiro, Assefa, B. Gapta, H. Ndue, M. Kissimou, Rene Nfon, Stuart Cable, Ben Pollard and the late Martin Etuge for assistance in the field. Drs Florence Ngo Ngwe, Eric Nana, Jean Betti Lagarde, the current and former directors, of IRAD-National Herbarium of Cameroon, Yaoundé, and their staff are thanked for expediting the collaboration between our two institutes. Janis Shillito typed the manuscript. Two anonymous reviewers are thanked for reviewing an earlier version of this paper.

